# Limb- and tendon-specific *Adamtsl2* deletion identifies a soft tissue mechanism modulating bone length

**DOI:** 10.1101/307496

**Authors:** Dirk Hubmacher, Stetson Thacker, Sheila M. Adams, David E. Birk, Ronen Schweitzer, Suneel S. Apte

## Abstract

Disproportionate distal limb shortening is the hallmark of acromelic dysplasias. Among them, geleophysic dysplasia is a rare, frequently lethal condition characterized by severe short stature, musculoskeletal, cardiac, pulmonary, and skin anomalies. Geleophysic dysplasia results from dominant fibrillin-1 (*FBN1*) or recessive *ADAMTSL2* mutations, suggesting a functional link between ADAMTSL2 and FBN1. Mice lacking ADAMTSL2 die at birth, precluding analysis of postnatal skeletal growth and mechanisms underlying the skeletal anomalies of geleophysic dysplasia. We show that *Adamtsl2* is expressed in limb soft tissues, predominantly in tendon. Expression in developing bones is limited to their terminal cell layers that are destined to become articular cartilage and is absent in growth plate cartilage. *Adamtsl2* conditional deletion in limb mesenchyme using *Prx1*-Cre led to an acromelic dysplasia, providing a suitable model for investigation of geleophysic dysplasia. Unexpectedly, conditional *Adamtsl2* deletion using *Scx*-Cre, a tendon-specific deleter, also impaired skeletal growth. Specific morphogenetic anomalies were seen in Achilles tendon, along with FBN1 accumulation. Thus, ADAMTSL2, shown here to bind fibrillin microfibrils in vitro, limits fibrillin microfibril formation in tendons and promotes tendon growth. The findings suggest that reduced bone growth in geleophysic dysplasia results from external tethering by short tendons rather than intrinsic growth plate anomalies.

## Introduction

Recessive *ADAMTSL2* mutations and dominant mutations located in exons 41-42 of *FBN1* cause geleophysic dysplasia (GD) in humans (MIM #231050 and MIM #614185, respectively) (1–3). GD is a rare skeletal disorder belonging to the acromelic dysplasia group and is characterized by short stature, joint contractures, tight skin, brachydactyly, delayed bone age, and hypermuscularity (4–8). Severe airway and cardiac involvement are common in GD and can result in death of affected children. A founder *ADAMTSL2* mutation in beagles causes Musladin-Lueke syndrome, which has a similar musculoskeletal phenotype as GD, but lacks lung or cardiac involvement and appears to run a milder course than GD (9, 10). In contrast to human GD, canine *FBN1* mutations causing Musladin-Lueke syndrome are not known.

ADAMTS-like 2 (ADAMTSL2) is a secreted glycoprotein belonging to the ADAMTS superfamily, which comprises 19 secreted metalloproteases (ADAMTS proteases) with diverse substrates, and 7 ADAMTS-like (ADAMTSL) proteins lacking a protease domain (11, 12). Thus, ADAMTSL proteins are thought to have non-proteolytic roles as extracellular matrix (ECM) components or to regulate ADAMTS protease activity as potential co-factors (13, 14). Recessive mutations in two ADAMTS proteases cause acromelic dysplasias, the resulting overlapping manifestations constituting a Weill-Marchesani syndrome (WMS) spectrum (*ADAMTS10*, MIM #277600; *ADAMTS17*, MIM #613195). WMS spectrum is distinguished from GD by several features, including predominant eye involvement which is not seen in GD, and it lacks the severe cardiopulmonary manifestations or juvenile lethality that is typical of GD (2, 15–18). In addition to phenocopying GD, dominant *FBN1* mutations also lead to WMS (MIM #608328) and another acromelic dysplasia, acromicric dysplasia (MIM #102370) (3, 19, 20). FBN1 is the major component of fibrillin microfibrils, which are 10-12 nm diameter fibrillar supramolecular structures in the ECM (21). Microfibrils provide structural integrity to tissues and regulate members of the transforming growth factor (TGF)-β superfamily by sequestering them directly (bone morphogenetic proteins) or via their binding to latent TGFβ binding proteins (22–24). The genetics of human acromelic dysplasias suggests that ADAMTS proteins and FBN1 operate in the same pathway to regulate musculoskeletal development and growth, and that ADAMTSL2 could affect the structure and function of microfibrils. However, the specific roles and mechanisms of ADAMTSL2 at the tissue and molecular level during musculoskeletal development remain to be fully elucidated.

*Adamtsl2* knockout mice die perinatally due to bronchial occlusion and a ventricular septal defect that precluded analysis of postnatal bone growth (25). Notably, skeletal patterning and development were not impaired in the null embryos, suggesting the possibility of a defective longitudinal bone growth during the juvenile period. We previously showed that recombinant ADAMTSL2 bound not only to FBN1, but also to FBN2 (25). FBN2, together with a third fibrillin isotype, FBN3 (present in humans but not in mice), is the embryonic fibrillin isotype, whereas FBN1 plays major roles in postnatal growth and tissue integrity (26–28). On the basis of these findings, it was suggested that ADAMTSL2 may modulate the balance between embryonic and adult fibrillin isotypes in microfibrils (13). Notably, In the *Adamtsl2* knockout mice, fibrillin-2 (FBN2) microfibrils accumulated at the interface of bronchial smooth muscle cells and the bronchial epithelium (25).

Here, we demonstrate that tissue-specific *Adamtsl2* deletion, either in limb mesenchyme or tendons, recapitulates the skeletal phenotype of GD. These experiments, together with elucidation of additional in vitro mechanisms, provide new insights on mechanisms of geleophysic dysplasia with broad relevance to skeletal growth regulation and tendon development.

## Results

### *Adamtsl2* mRNA is expressed in musculoskeletal soft tissues and presumptive articular cartilage, but not growth plate cartilage

The temporal and spatial pattern of *Adamtsl2* mRNA expression during embryonic mouse musculoskeletal development and during postnatal growth was revealed by β-galactosidase staining using an intragenic *Adamtsl2*-lacZ reporter transgene, which provided a surrogate for *Adamtsl2* mRNA expression. At embryo age 14.5 (E14.5) and E17.5 whole limb staining showed that *Adamtsl2* was most strongly expressed in developing tendons (Figure 1A, B). Stained sections showed strong *Adamtsl2* expression in superficial layers of the presumptive articular cartilage (Figure 1C, D) and in a subset of skeletal muscle cells forming muscle spindles (Figure 1E, F). At postnatal day 12 (P12), *Adamtsl2* was strongly expressed in tendons throughout the limbs, including the Achilles tendon (Figure 1G, H). In the skin, *Adamtsl2* was expressed in a thin layer of cells above the panniculus carnosus (Figure 1I, arrows). In skeletal muscle, regions of intense, localized *Adamtsl2* expression were observed, attributed to muscle spindles (Figure 1I, J, arrows in J). In bones and joints, *Adamtsl2* expression continued in the superficial layer of the articular cartilage, but was consistently absent from the growth plate and bone (Figure 1K-M). In the knee joint, *Adamtsl2* was also expressed in superficial cells of the meniscus (Figure 1L arrow heads), in addition to the articular cartilage zones of the ends of the tibia, fibula and femur (Figure 1L, arrows). In the vertebral column, we detected strong *Adamtsl2* expression in the inner annulus fibrosus layer of the intervertebral disc, but not in the nucleus pulposus (Figure 1N, Q).

**Figure 1.**
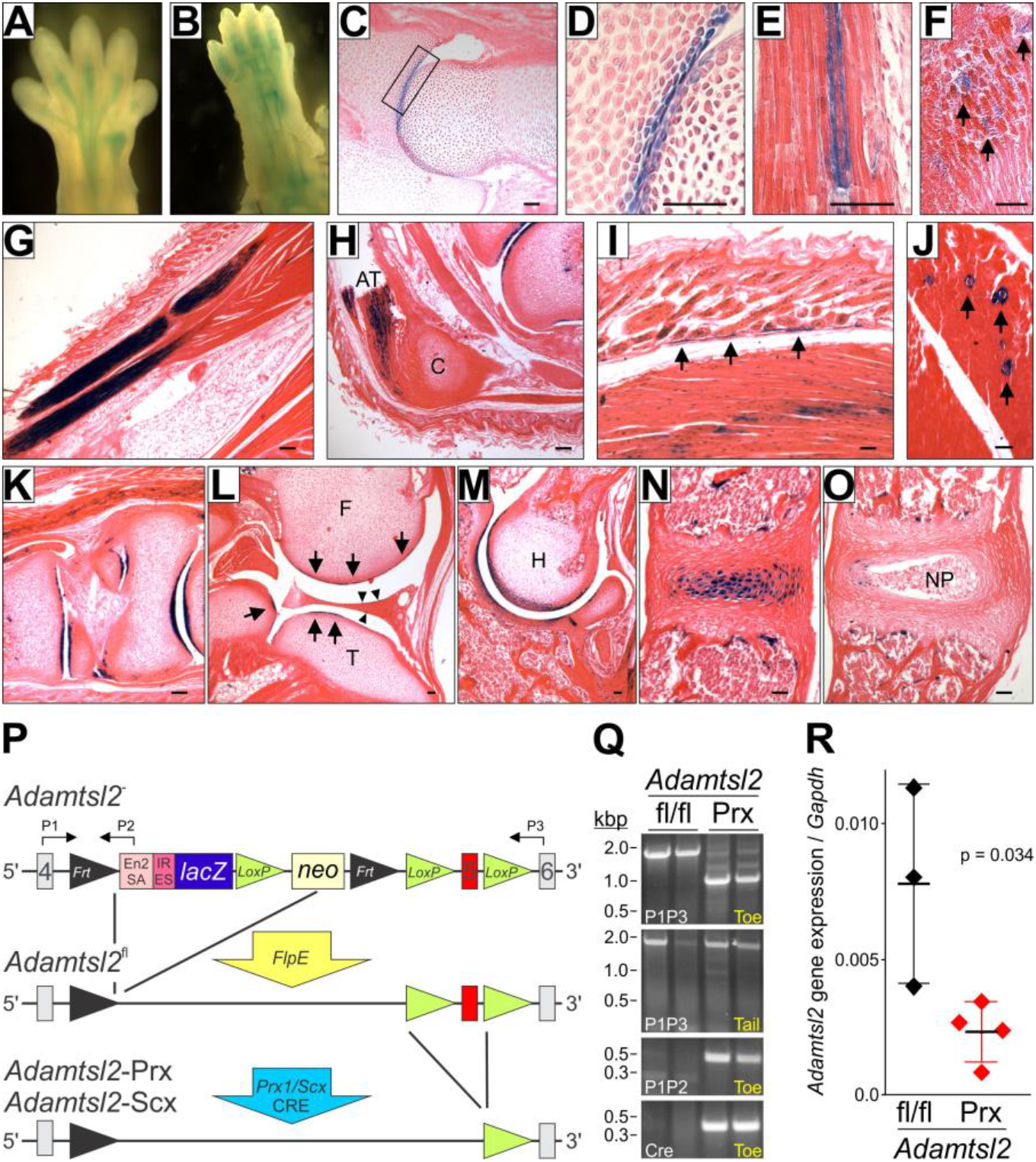
*Adamtsl2* expression in specific musculoskeletal tissues and conditional deletion in limb mesenchyme and tendon. (**A, B**) After whole limb β-gal staining, *Adamtsl2* expression (blue) was observed in developing forelimb tendon at E14.5 (A) and E17.5 (B). (**C-F**) At E17.5, *Adamtsl2* was expressed in prospective articular cartilage (C, D; proximal femur. Panel D represents higher magnification of boxed area in C), and skeletal muscle (E, F). Arrows in panel F indicate focal expression in muscle. (**G-O**) At P12, *Adamtsl2* was expressed in tendons (G, H), dermis (I; arrows indicate panniculus carnosus), skeletal muscle (I, J), prospective articular cartilage in the hind foot (K) and knee joints (L; arrows indicate joint lining cells, arrowheads indicate outer meniscal cell layer), shoulder joint (M), and the inner annulus fibrosus layer, but not the nucleus pulposus of the intervertebral disc (N, O; N is a tangential section that does not contain the nucleus pulposus). Images A-O are representative of n=2-3. Scale bars in A-O are 50 μm. AT, Achilles tendon, C, calcaneus; F, femur; H, humerus; NP, nucleus pulposus; T, tibia. (**P**) Gene targeting strategy for limb- and tendon-specific *Adamtsl2* deletion. The *lacZ-Δ*neo cassette was removed with *FlpE* (*Adamtsl2*^fl^) and *Adamtsl2* was inactivated by excision of exon 5 with *Prx1*-Cre or *Scx*-Cre (*Adamtsl2*-Prx, *Adamtsl2*-Scx, respectively). (**Q**) PCR from toe and tail genomic DNA (gDNA). The band shift with primer pair P1-P3 from toe but not tail gDNA indicates excision of exon 5. (**R**) RT-qPCR of Achilles tendon RNA shows reduction in *Adamtsl2* expression after gene deletion (n=3). P-value was calculated with a 2-sided Student t-test.

### Conditional *Adamtsl2* deletion in limb-mesenchyme leads to acromelic dysplasia

To define the role of ADAMTSL2 in post-natal limb growth, we bypassed neonatal lethality of the *Adamtsl2* null mice by using *Prx1*-Cre-mediated limb mesenchyme-specific inactivation of *Adamtsl2* (Figure 1P) (25, 29). PCR analysis of genomic DNA extracted from toes, but not from tail DNA indicated successful limb-specific excision of the floxed exon-5 of *Adamtsl2* by Cre-recombinase (genotype referred to as *Adamtsl2*-Prx) (Figure 1Q). In addition, real-time quantitative PCR (RT-qPCR) of Achilles tendon mRNA showed a significant reduction of *Adamtsl2* mRNA in *Adamtsl2*-Prx mice compared to control tendons (Figure 1R). *Adamtsl2*-Prx mice were viable and without gross musculoskeletal anomalies.

Alizarin red-Alcian blue stained skeleton preparations from P16 to P20 demonstrated that *Adamtsl2*-Prx long bone diaphyses were less sculpted, i.e., they lacked a characteristic narrowing in the central diaphyseal regions and had a “stubby” appearance (2A, B, F-H, “D” indicates narrowest part of diaphyses). These changes were most visible in metacarpals and metatarsals (Figure 2B, H). All forelimb bones were reduced in length in *Adamtsl2*-Prx mice (Figure 2C, D). The distal bones were disproportionately shorter, indicating an acromelic limb phenotype. The diaphyseal and metaphyseal regions of *Adamtsl2*-Prx forelimb bones were significantly wider compared to controls (Figure 2E). Like the forelimbs, *Adamtsl2*-Prx hindlimb bones were also shorter and their diaphyseal and metaphyseal regions were significantly wider in *Adamtsl2*-Prx limbs (Figure 2F-K). At birth (P0), metatarsals and radii of *Adamtsl2*-Prx mice were significantly shorter, but no significant differences were observed in the length of metacarpals and the ulna (Supplemental Figure 1A). In *Adamtsl2*-Prx mice older than 9 months, significant bone shortening persisted in metatarsals and the tibia, but was not observed in the forelimb (metacarpals, humerus, radius, ulna) (Supplemental Figure 1B). Neither histological abnormalities in the growth plate of *Adamtsl2*-Prx limbs nor differences in the height of the growth plate, or its proliferative, pre-hypertrophic and hypertrophic zones were detected (Supplemental Figure 2A, B). In summary, limb-specific deletion of *Adamtsl2* consistently resulted in distal bone shortening in the limbs and altered the shape of the diaphyseal and metaphyseal regions, i.e., recapitulating skeletal anomalies described in GD.

**Figure 2.**
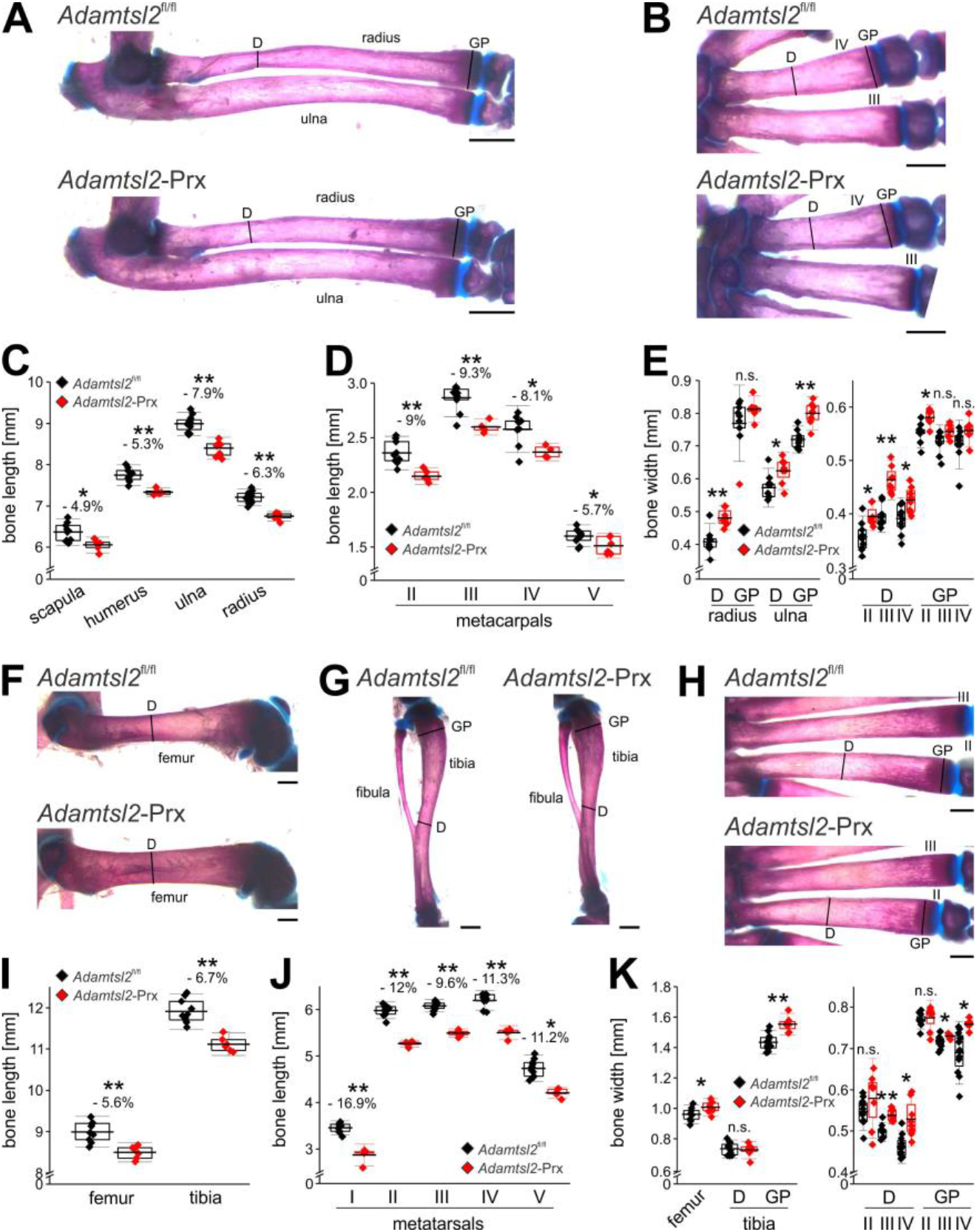
*Prxl*-Cre mediated *Adamtsl2* inactivation results in an acromelic dysplasia (disproportionally shorter distal bones). **(A, B)** Alizarin red-Alcian blue stained radius and ulna (A) and metacarpals (B), showing the sculpting defect in the mid-diaphysis. Lines indicate where bone width was measured. D, diaphysis; GP, growth plate. **(C, D)** Measured lengths of proximal forelimb long bones (C) and metacarpals (D) indicating disproportionate shortening of the distal bones. (**E**) Measured widths of diaphysis (D) or growth plate (GP) indicate wider bones in absence of ADAMTSL2. Relative reduction in length or width in C-E is indicated as a percentage. (**F-H**) Alizarin red-Alcian blue stained femur (F), tibia and fibula (G), and metatarsals (H), showing the sculpting defect in the diaphysis. Lines indicate where bone width was measured. (**I, J**) Measured lengths of hindlimb proximal long bones (I) and metatarsals (J) indicating disproportionately shorter distal bones. (**K**) Measured widths of diaphysis (D) or growth plate (GP) indicating wider bones in absence of ADAMTSL2. Relative reduction in length or width in I-K is indicated as a percentage. Bones were dissected from littermates at P18 (n=6-10). Relative reduction in length is indicated in %. P-value were calculated with 2-sided Student t-test. * p<0.05; ** p<0.001. The box indicates the 25^th^-75^th^ percentile, line indicates mean value and whiskers indicate standard deviation. Scale bars in A-C and F-H are 1 mm.

### Abnormal Achilles tendon in *Adamtsl2*-Prx mice

In addition to the skeletal anomalies, Achilles tendons from *Adamtsl2*-Prx mice had altered dimensions and morphology (Figure 3A, B). Specifically, their origins from the gastrocnemius were poorly defined, and they were significantly shorter and wider than controls (Figure 3A, B). During limb dissection, we also noticed that *Adamtsl2*-Prx Achilles tendons were tethered to surrounding tissue, reminiscent of severe peri tendon fibrosis reported in Achilles tendon from dogs with Musladin-Lueke syndrome (10). In contrast, the control Achilles tendons could be readily separated from the surrounding tissue. Because tendons consist predominantly of collagen I fibrils and genetic loss of specific ECM components alters collagen fibril diameter (30–33), we analyzed the cross-sectional diameter of the collagen fibrils by transmission electron microcopy. The profiles of the collagen fibrils were regular and comparable in the two groups. Based on a two-sample Kolmogorov-Smirnov test, no significant difference in the distribution of the collagen fibril diameter was detected in Achilles tendon tissues from *Adamtsl2*-Prx limbs compared to control tendons (Figure 3C, D).

**Figure 3.**
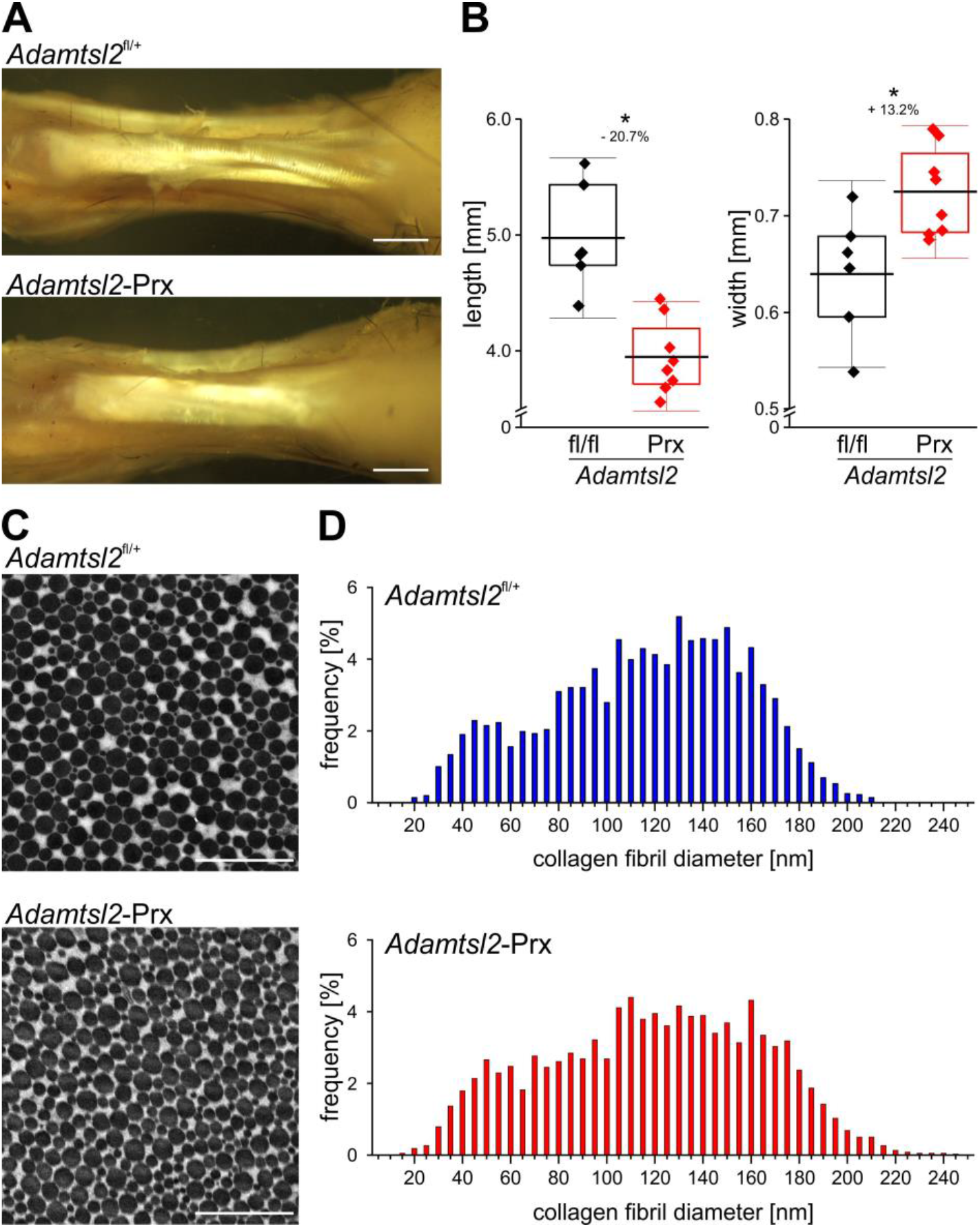
Altered Achilles tendon morphology in the absence of *Adamtsl2*. **(A)** A dissection of Achilles tendon at P27 shows reduced length and altered morphology of ADAMTSL2-deficient tendon (increased width, poorer definition of sub-tendons at their origin from the gastrocnemius muscle (right)). Images shown are representative of n=6-8 and represent littermates. Scale bars are 1 mm. (**B**) *Adamtsl2*-deficient Achilles tendons are significantly shorter and wider than wild-type (n=6-8). P-values, calculated with a two-sided Student t-test and relative reduction in length or increase in width in *%* are indicated. * p<0.05. The box indicates the 25^th^-75^th^ percentile, line indicates mean value and whiskers indicate standard deviation. **(C)** Electron micrograph of crosssection through the Achilles tendon mid-substance showing that mutant and wild-type collagen fibrils have comparable cross-sectional morphology. Images shown are representative of n=10-12 images and n=3 littermates per genotype. Scale bars represent 1 μm. **(D)** Histograms of the distribution of the collagen fibril diameters indicated comparable distribution profile between the genotypes (10-12 digital images were quantified from n=3 mice per genotype). No significant difference between the distribution of the two data sets was detected in a Kolmogorov-Smirnov test (D=0.1569, p=0.521).

Given intense *Adamtsl2* expression in tendons and pronounced morphological alterations in *Adamtsl2*-Prx Achilles tendons, we undertook tendon-specific *Adamtsl2* deletion using *Scx*-Cre (*Adamtsl2*-Scx). The mT/mG reporter mouse, which switches cellular fluorescence from red to green in the presence of Cre demonstrated deletion specifically in tendons, including the Achilles tendon but not the distal region of long bones nor skeletal muscle (Figure 4A-D). Although Achilles tendons of *Adamtsl2*-Scx limbs were significantly shorter, their visually increased width did not reach statistical significance over control tendons (Figure 4E, F). As in *Adamtsl2*-Prx limbs, several *Adamtsl2*-Scx forelimb and hindlimb bones were shorter, with a ~6% reduction in length (Figure 4G, H), but disproportionate shortening of distal bones found in *Adamtsl2*-Prx limbs was not seen in *Adamtsl2*-Scx limbs. The length of metacarpals from *Adamtsl2*-Scx forelimbs was not different compared to control limbs (data not shown). Thus, deletion of *Adamtsl2* in tendons led to substantially similar but milder changes than those arising from *Adamtsl2* deletion in total limb mesenchyme.

**Figure 4.**
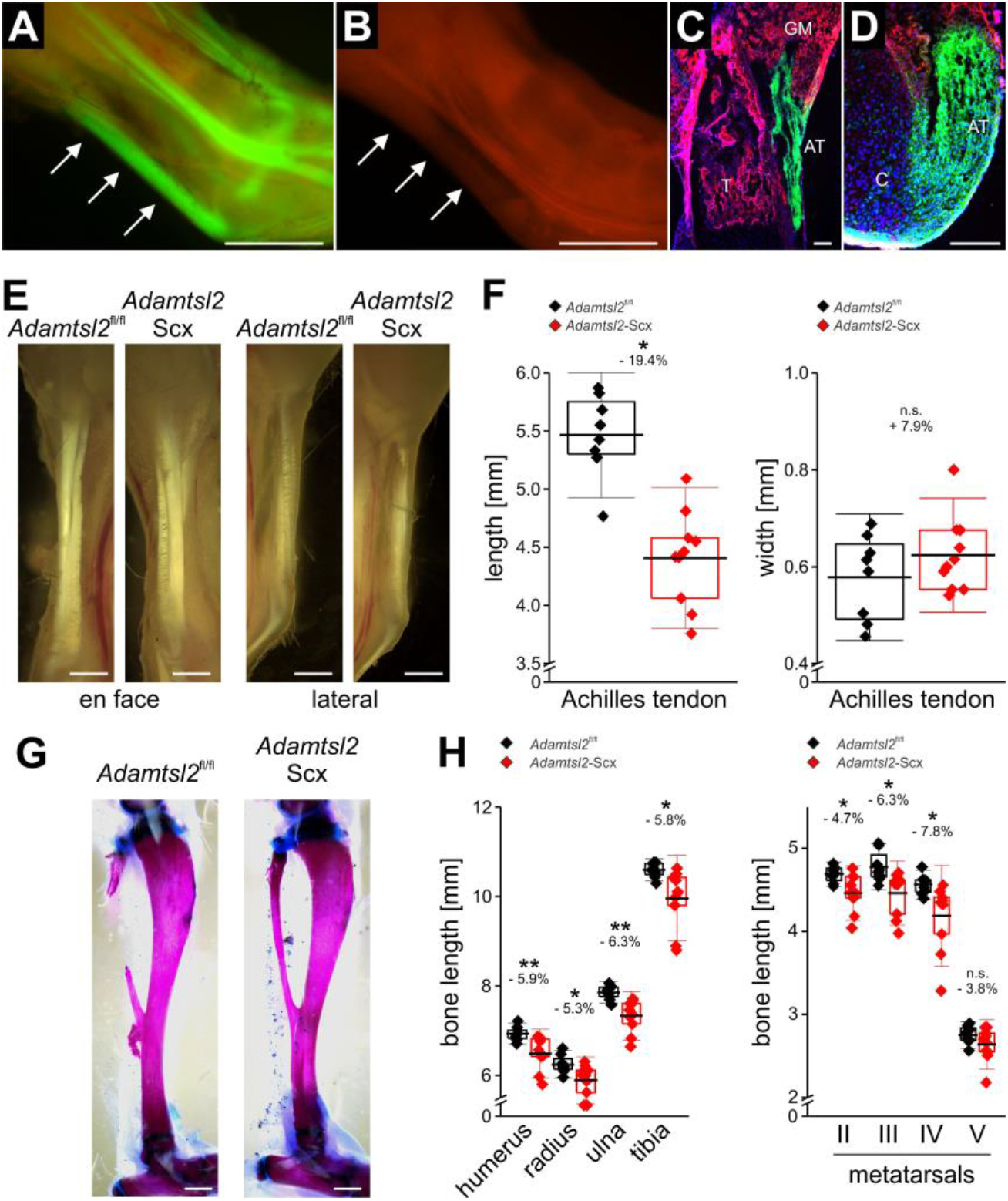
*Scx*-Cre mediated *Adamtsl2* inactivation in tendon results in altered tendon morphology and non-autonomous bone shortening. **(A, B)** Fluorescence imaging of the ankle from newborn *Adamtsl2-Scx* mice shows green fluorescence in tendons indicating specific *Scx*-Cre recombinase activity (A, arrows indicating Achilles tendon) compared to red fluorescence in *mG/mT* mice without *Scx*-Cre (B, arrows). Scale bars in A and B are 1 mm. Images shown are representative of n=2 littermates. **(C, D)** Longitudinal section through distal tibia (T) showing *Scx*-Cre activity in Achilles tendon tissue (AT, green), but neither gastrocnemius muscle (GM), tibial bone (red) (C) nor the calcaneal insertion (C) of the Achilles tendon showed *Scx*-Cre activity. (D). Scale bars in C and D represent 100 μm. **(E)** Gross view of Achilles tendon shows altered dimensions from the dorsal (left-hand panels) or lateral aspects (right-hand panels). Images shown are representative of n=8-10 from n=4-5 littermates. Scale bars in E are 1 mm. **(F)** *Adamtsl2*-deficient Achilles tendons are significantly shorter and tended to be wider (n=8-10). Relative reduction in length is indicated in %. **(G)** Alizarin red-Alcian blue stained tibiae showing normal morphology. Scale bars represent 1 mm. **(H)** Bone length measurements of forelimb and hind limb long bones and metatarsals demonstrates bone shortening in *Adamtsl2*-Scx limbs (n=8-10). Relative reduction in length is indicated in %. P-values were calculated with 2-sided Student t-test. * p<0.05; ** p<0.01, n.s. not significant. The box indicates the 25^th^-75^th^ percentile, the line indicates mean value and whiskers indicate standard deviation.

### ADAMTSL2 deficiency alters tenocyte organization

Longitudinal sections through the tendon mid-substance revealed a disorganization of tenocytes in *Adamtsl2*-Prx tendons (Figure 5A, B). In control tendons, elongated tenocytes form regular linear arrays oriented along the longitudinal axis of the tendon. However, in *Adamtsl2*-Prx tendon, the tenocytes were rounded with loss of register in tenocyte arrays. Multiphoton imaging (second harmonic generation) demonstrated a disarray of collagen fiber orientation and higher fluorescence intensity (Figure 5C, D). *Adamtsl2* deficient Achilles tendon ECM had more intense FBN1 staining in the pericellular matrix around the tenocyte arrays than control tendons (Figure 5E, F). FBN2 staining intensity was low and demonstrated no obvious difference between *Adamtsl2* deficient and control tendons (data not shown). *Fbn1*, *Fbn2*, *Fn1*, *Col1a1*, *Col3a1*, *Scx*, and *Tnmd* mRNA levels were not altered in *Adamtsl2*-Prx tendons (Supplemental Figure 3).

**Figure 5.**
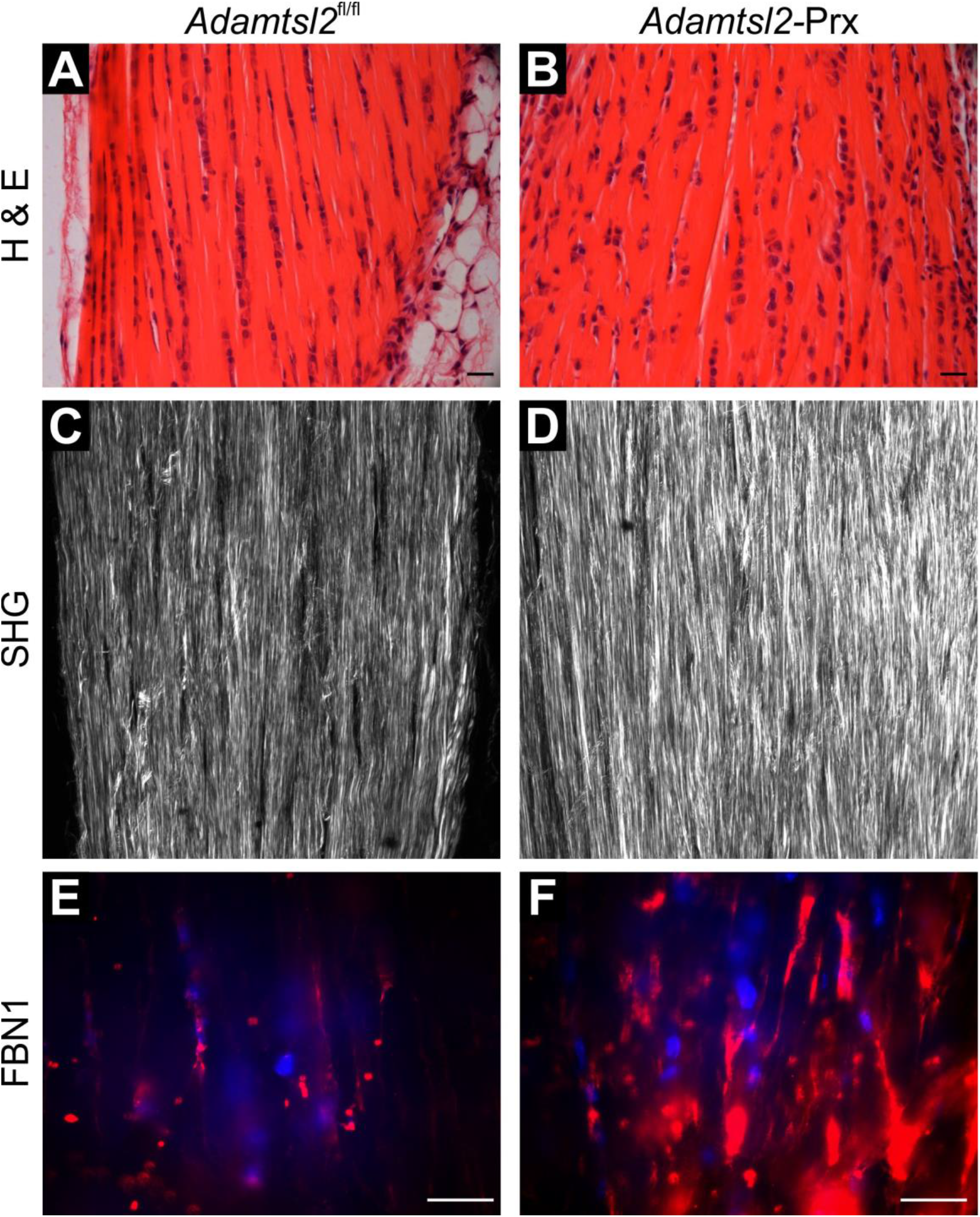
Disorganization of tenocyte arrays and pericellular matrix in *Adamtsl2*-Prx tendon. **(A, B)** Hematoxylin & eosin staining of longitudinal sections through Achilles tendon show disturbance of linear tenocyte arrays in *Adamtsl2*-Prx tendon. Images shown are representative of n=3 littermates per genotype. **(C, D)** Second harmonic generation imaging of collagen fibers shows more intense collagen signal and disorganized collagen fibers in the absence of ADAMTSL2. Images shown are representative of n=3 littermates. **(E, F)** Fibrillin-1 (FBN1) staining is cell-associated, and stronger in *Adamtsl2*-Prx tendons. Images shown are representative of n=3 littermates. Scale bars represent 20 μm.

We showed previously that recombinant ADAMTSL2 directly bound to the N- and C-terminal halves of recombinant FBN1 and FBN2 (3, 25). To discern whether ADAMTSL2 also bound to the corresponding supramolecular complexes, i.e., to fibrillin microfibrils, we added purified recombinant ADAMTSL2 to human dermal fibroblasts and asked whether it colocalized with endogenous fibrillin microfibrils. Recombinant ADAMTSL2 co-localized with both FBN1 and FBN2 microfibrils, but the colocalization with FBN2 was weaker because fewer FBN2-stained microfibrils were formed by these cells (Figure 6A). To capture early time points of microfibril formation and to analyze the potential co-localization of ADAMTSL2 with fibronectin fibrils, whose assembly precedes that of fibrillin microfibrils, we performed co-immunostaining experiments 24h after cell seeding and recombinant ADAMTSL2 supplementation (Figure 6B). We found that ADAMTSL2 co-localized with nascent FBN1 microfibrils, but not with FBN2 at the 24h time point. ADAMTSL2 also co-localized to some, but not all, fibronectin-positive fibrils. Given the previously published co-localization of FBN1 or FBN2 with fibronectin (34) and colocalization of FBN1 and FBN2 (35, 36), our data suggest that ADAMTSL2 predominantly binds to FBN1 microfibrils in the ECM of human dermal fibroblasts. Taken together with increased FBN1 immunostaining in *Adamtsl2*-Prx tendons, the data suggest a specific regulatory role for ADAMTSL2 in limiting FBN1 levels in tendon microfibrils after birth.

**Figure 6.**
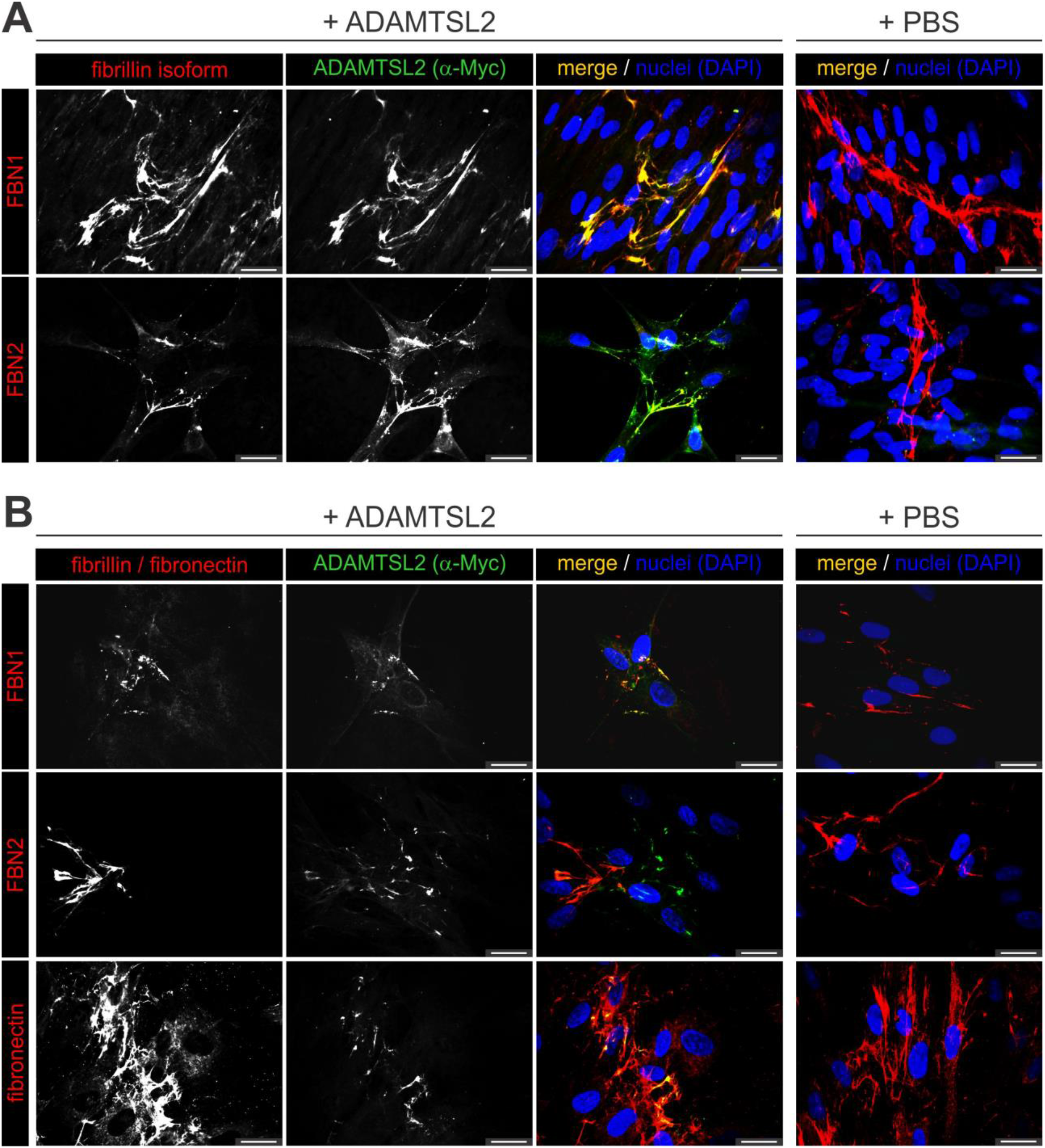
ADAMTSL2 co-localizes with fibrillin microfibrils in cultured fibroblasts. **(A)**ADAMTSL2 co-localizes with fibrillin-1 (FBN1) and to a lesser extent with fibrillin-2 (FBN2) containing microfibrils assembled by human dermal fibroblasts 48h after seeding cells and addition of recombinant ADAMTSL2. **(B)** 24h after seeding cells and ADAMTSL2 addition, ADAMTSL2 localized to FBN1 microfibrils and fibronectin fibers. No localization to FBN2 microfibrils was observed. Nuclei were counterstained with DAPI. Scale bars are 20 μm.

## Discussion

Here, we report generation and analysis of mouse models that recapitulate skeletal manifestations of human geleophysic dysplasia. Our studies identified an unexpected distribution of *Adamtsl2* mRNA, showing its absence in cartilage growth plates, where skeletal growth occurs, but intense expression in tendons. Tendon-specific *Adamtsl2* deletion suggested that ADAMTSL2 expressed by tendons and by implication, tendons themselves, non-autonomously regulate skeletal growth. Intense *Adamtsl2* expression in tendons, tenocyte disarray, ADAMTSL2 binding to fibrillin microfibrils and excess FBN1 microfibrils in tendons upon ADAMTSL2 deletion strongly suggests that ADAMTSL2 limits formation of fibrillin microfibrils in tendon pericellular matrix, and has a tissue-autonomous function in tendon.

Bone growth results from chondrocyte proliferation, hypertrophy and ECM synthesis at cartilage growth plates, located at the ends of long bones. The mediators of growth are well-characterized and include Indian hedgehog and parathyroid hormone-related protein acting locally in the growth plate, and endocrine factors such as growth hormone, and vitamin D (37–39). Soft tissues have long been known to have a role in bone growth, surmised from limb shortening after soft tissue contractures, juvenile paralysis or experimental manipulation of soft tissues (40–42). The skeletal impact of soft tissue anomalies is thought to arise from mechanical constraints imposed by reduced soft tissue extensibility, consistent with retardation of growth plate activity by exogenous compressive force (43, 44). Notably, all tendons cross at least one joint, allowing the muscle-tendon unit to exert compressive force on one or more growth plates. The lack of *Adamtsl2* expression in growth plate cartilage suggests that a local, direct role on intrinsic cartilage growth control pathways is unlikely. In contrast, extremely strong *Adamtsl2* expression in tendons, shortening of tendons observed after both *Prx1*-Cre and *Scx*-Cre-mediated conditional deletion, and clinically manifest peri-tendon fibrosis and contractures in patients with GD and dogs with Musladin-Lueke syndrome (9, 10, 45) support a specific role for ADAMTSL2 in tendon growth and a growth plate-extrinsic mechanism affecting long bone growth. Moreover, tendon-specific deletion with *Scx*-driven Cre, which, unlike *Prx*-Cre, spares recombination in cartilage, specifically led to bone shortening. The present studies have not formally excluded a potential biochemical regulatory pathway in which products released from tenocytes would act in an endocrine / paracrine fashion on the growth plate.

Previous work showed that linear tenocyte arrays in maturing tendons were specifically associated with fibrillin microfibrils, which were postulated to represent fiducial elements essential for tenogenesis (46). Although our study showed that collagen fibrillogenesis was unimpaired, the findings suggest that accumulation of FBN1 in tendon microfibrils affects the overall growth and morphogenesis of the Achilles tendon, including the alignment and abundance of collagen fibers, which are higher-order collagen assemblies of fibrils. The observed minor differences in skeletal growth following *Prx1*-Cre and *Scx*-Cre mutagenesis may arise from different temporal and spatial deletion by these promoters. ADAMTSL2 deletion in muscle and skin by *Prx1*-Cre, where it was not deleted by *Scx*-Cre, may also contribute an additional impact on skeletal growth. Interestingly, *Fbn1* deletion using *Scx*-Cre results in bone lengthening i.e., a mouse model for skeletal manifestations of Marfan syndrome (Dr. Francesco Ramirez, Icahn School of Medicine at Mount Sinai, personal communication). What is presently unclear is the genesis of the cortical diaphyseal and metaphyseal bone sculpting defect that was observed, since ADAMTSL2 is not expressed in bone of periosteum. We suggest that this could arise from the influence of contracted tendons and soft tissue in the juvenile period prior, since bone remodeling is sensitive to mechanical forces (47).

At the molecular level, the prior demonstration that ADAMTSL2 binds to both FBN1 and FBN2, is extended here to demonstrate their binding to microfibrils which in addition to fibrillins, contain numerous other molecules (1, 25). Previously, in embryos with global loss of ADAMTSL2, we identified an excess of FBN2 microfibrils in bronchial ECM associated with bronchial epithelial dysplasia and occlusion of bronchial lumena (25). FBN2 is the major fibrillin expressed during embryogenesis, with FBN1 expressed at relatively low levels until the end of gestation (26, 27). At birth, which represents a watershed in fibrillin gene expression, FBN2 expression wanes and FBN1 becomes the dominant fibrillin isotype in mice. Thus, in addition to FBN2 accumulation in embryonic *Adamtsl2*-/- bronchi, intense FBN1 staining was seen in postnatal *Adamtsl2*-deficient Achilles tendon. These findings, taken together with the observation that ADAMTSL2 has at least two binding sites on both fibrillin isotypes (1, 48), suggest a general function for ADAMTSL2 in limiting microfibril assembly, i.e., ADAMTSL2 may suppress microfibril formation independent of fibrillin isotype (3, 25). However, when recombinant ADAMTSL2 was added to human dermal fibroblasts, no consistent change in the amount of fibrillin microfibril staining was observed. One possibility is that dermal fibroblasts do not recapitulate the precise expression of microfibril proteins and ADAMTS proteins as tenocytes. Alternatively, ADAMTSL2 may modulate protease activity in the ECM, interfering with microfibril turnover in tissues. Since FBN2 and FBN1 microfibrils accumulated in the absence of ADAMTSL2 in vivo, ADAMTSL2 may act as a positive regulator of protease activity against microfibrils in vivo. In this regard, mutation of two ADAMTS proteases, ADAMTS10 and ADAMTS17 results in acromelic dysplasias, and ADAMTS10 has limited proteolytic activity against fibrillin-1 (49). These mechanisms have not been tested in cultured tenocytes, which do not retain the specific tenocyte phenotype in culture.

The enduring paradox of *FBN1* mutations is that the majority cause MFS, but a subset of mutations, located in the fifth transforming growth factor-β-like / 8-cysteine domain of FBN1 (exons 41 and 42), lead to acromelic dysplasias, whose manifestations are the opposite of MFS. Jensen et al demonstrated that MFS-causing *FBN1* mutations affected molecular secretion, and could thus have a haploinsufficiency effect, whereas *FBN1* mutants causing GD, which affect a specific domain, TB5, were secreted and assembled into microfibrils (50). TB5 mutants leading to GD block the binding of this domain to heparan-sulfate, which regulates microfibril assembly (51). Our findings regarding ADAMTSL2-fibrillin microfibril interactions on the one hand, and the effect of TB5 of mutations on the other, would support a role for a specific interaction between ADAMTSL2 and TB5 that inhibits microfibril assembly.

In summary, we present a novel mouse model for acromelic dysplasia recapitulating skeletal anomalies of GD. We discovered that shortening of the Achilles tendon in the absence of ADAMTSL2 influences bone growth, presumably by creating a biomechanical restraint on bone growth. An important direction for future investigation is how ADAMTSL2 regulates tenocyte alignment and influences tendon growth and collagen assembly.

## Methods

All reagents were purchased from Sigma-Aldrich or ThermoFisher Scientific unless specified.

### Mouse strains

*Adamtsl2-Prx* mice were generated by deleting the Frt-site-flanked *lacZ*-Δneo cassette from *Adamtsl2*^+/-^ mice (KOMP allele: *Adamtsl2*^tm1a(KOMP)Wtsi^, NIH, Bethesda, MD, USA) using B6.Cg-Tg(*ACTB-FLPe*)9205Dym/J (Jackson Laboratory, Bar Harbor, ME USA) (25). FLPe (FLP recombinase)-mediated deletion generated the *Adamtsl2*^fl^ allele, where exon 5 is flanked by loxP sites and is amenable to Cre-mediated excision (Figure 1D). A *B6.Cg-Tg(Prrx1-cre)1Cjt/J* (*Prx1*-Cre) male mouse was obtained from Jackson Laboratory and the *Scx*-Cre strain was previously described (52).

### Genotyping

Genomic DNA from toe and tail tissue was isolated using DirectPCR (Viagen, Los Angeles, CA, USA). PCR products were amplified using *Taq* polymerase (New England BioLabs Inc., Ipswich, MA, USA) and a forward primer located in exon 4 (*Adamtsl2* wild-type allele, P1: 5´-gtaccagctctgcagagtgc-3´) in combination with reverse primers located in the *En2*-splice acceptor site in the *lacZ*-Δneo (*Adamtsl2* knock-out allele, P2: 5´-cactgagtctctggcatctc-3´) or in exon 6 (*Adamtsl2* wild-type or conditional allele, respectively, P3: 5´-ctctcaggtcggtgagcttg-3´). PCR products were separated by agarose gel electrophoresis and visualized with ethidium bromide.

### β-Galactosidase (β-gal) staining

Tissue was fixed in 4% paraformaldehyde (Electron Microscopy Sciences, Hatfield, PA, USA) overnight and stained with potassium ferrocyanide/potassium ferricyanide/5-bromo-4-chloro-3-indolyl-β-D-galactopyranoside (X-gal) (Denville Scientific Inc., Holliston, MA, USA) as described previously (53).

### Bone and tendon morphometry

Limbs were dissected, and the Achilles tendon exposed and photographed in situ. The soft tissue was removed, and bones were fixed in 80% ethanol and dehydrated in 96% ethanol and acetone over several days for Alizarin red / Alcian blue staining. Staining was performed with a solution of 30 mg Alcian blue and 5 mg Alizarin red in 20 ml acetic acid / 80 ml 95% ethanol for several days at room temperature. Tissues were rinsed with 95% ethanol and cleared with 1% aqueous potassium hydroxide followed by serial transfer in 20%, 50% and 80% glycerin prepared in 1% aqueous potassium hydroxide. Cleared and stained bones were photographed, and bone length was measured using Image J (NIH, Bethesda, MD). Mutants were compared to littermate controls (*Adamtsl2*^fl/fl^, *Prx1*-Cre, *Scx*-Cre) in all analyses and statistical significance was determined with a two-sided Student t-test using the Origin 2017 software package (OriginLab, Northampton, MA, USA).

### Gene expression analysis

Achilles tendons were homogenized with an Ultra Turrax homogenizer (IKA-Works Inc., Wilmington, NC, USA) in Trizol and total RNA was extracted according to the manufacturer’s protocol. 1 μg of total RNA was reverse transcribed using the High Efficiency cDNA Reverse Transcription kit (Applied Biosystems, Foster City, CA, USA). Quantitative realtime PCR (qPCR) was performed with EvaGreen (VWR, Radnor, PA, USA) in a total volume of 10 μl using 0.125 μl cDNA template. qPCR reactions were performed in triplicate with a CFX96 real-time system (Bio-Rad, Hercules, CA, USA).

### Electron microscopy

All chemicals for electron microscopy were purchased from Electron Microscopy Sciences (Hatfield, PA, USA). Achilles tendons were fixed immediately upon dissection and then overnight in cold 4% paraformaldehyde, 2.5% glutaraldehyde, 0.1 M sodium cacodylate and 8 mM CaCl_2_, pH 7.4, rinsed in cacodylate buffer and post-fixed for 1 h with 1% osmium tetroxide. Fixed tendons were dehydrated in a graded ethanol series followed by 100% propylene oxide, infiltrated and embedded over a 3-day period in a mixture of Embed 812, acidic methyl anhydride, dodecenylsuccinic anhydride and 2,4,6-Tris(dimethylaminomethyl)phenol (DMP-30) and polymerized overnight at 60 °C. Cross-sections of tendons (90 nm) were prepared using a Leica ultramicrotome (Leica Microsystems Inc., Buffalo Grove, IL, USA) and post-stained with 2% aqueous uranyl acetate and 1% phosphotungstic acid, pH 3.2. The sections were examined and imaged at 80 kV using a JEOL 1400 transmission electron microscope (JEOL Ltd., Tokyo, Japan) equipped with a Gatan Orius widefield side mount CC Digital camera (Gatan Inc., Pleasanton, CA, USA). Tendon diameter analysis was obtained from pooled data from one tendon from each of three littermates of each genotype. Digital images from each tendon were taken from non-overlapping areas at 60,000X. 10-12 images were randomized and masked before fibril diameters were measured using a RM Biometrics-Bioquant Image Analysis System (Nashville, TN, USA). 3586 and 3797 fibril diameters for the control and the *Adamtsl2*-Prx genotype, respectively, were measured. Fibril diameters were measured along the minor axis of the fibril cross-section. Tendon diameter measurements were pooled into groups by genotype and represented as histograms. A Kolmogorow-Smirnow test was performed to determine if the two datasets were different (http://www.physics.csbsju.edu/stats/KS-test.html).

### Histology and immunostaining

Limbs were fixed in 4% paraformaldehyde (Electron Microscopy Sciences) in PBS for 24h. Those containing mineralized bone were subsequently decalcified in 14% EDTA solution, changed every 3 days for 3-4 weeks and paraffin-embedded. 8-10 μm sections were used for hematoxylin and eosin staining and Masson trichrome stain according to standard protocols. For immunostaining with FBN1 and FBN2 antibodies, tissues were rehydrated, and antigen retrieval was performed with citrate-EDTA for 4 x 1.5 min in a microwave oven. Sections were cooled in antigen-retrieval buffer rinsed with water and 2 x PBS and blocked for 1h in 5% normal goat serum in PBS. Sections were incubated with FBN1 antibody (54) diluted in blocking buffer overnight at 4 °C, rinsed 3 × 10 min in PBS, and incubated with secondary antibody (goat anti-rabbit-Alexa-564) (Jackson ImmunoResearch, West Grove, PA, USA) diluted in blocking buffer for 1h at RT. Sections were rinsed 3 × 10 min in PBS and mounted using VectaShield Gold with DAPI. Sections were photographed on an Olympus BX51 upright microscope (Olympus, Center Valley, PA, USA) using a Leica DFC7000T camera and Leica Application Suite v4.6 imaging software (Leica Microsystems, Wetzlar, Germany).

### Second-harmonic generation / Two-photon imaging

Paraformaldehyde-fixed, paraffin-embedded tendon sections were rehydrated and coverslipped in 50% glycerol in PBS. Images were acquired using a Leica TCS SP5 II Confocal/Multi-Photon high-speed upright microscope with a 25X water immersion lens in forward scattering mode (Leica Microsystems). The excitation wavelength was set to 880 nm and the non-descanned detector (NDD) was set to collect signals between 430 and 450 nm. Z-stacks were acquired at a scan speed of 600 Hz in bidirectional mode with a line average of 5. Images were processed using ImageJ-Fiji software (NIH, Bethesda, MD).

### Cell culture and co-localization studies

Human dermal fibroblasts derived from explant cultures of circumcised foreskin were cultured in Dulbecco’s Modified Eagle Medium (DMEM) supplemented with 100 units/ml penicillin, 100 μg/ml streptomycin, and 5 mM L-glutamine (complete DMEM) in a 5% CO_2_ atmosphere in a humidified incubator at 37 °C. 50,000 cells / chamber were seeded in 8-well chamber slides (BD Bioscience, San Jose, CA, USA) and after 16 h 50 μg of recombinant mouse ADAMTSL2 (12), was added in complete DMEM (25). After 24 h, 48 h, or 72 h cells were fixed in ice-cold 70% acetone / 30% methanol (v/v) and co-immunostained for recombinant ADAMTSL2 using monoclonal mouse α-myc antibody (clone 9E10, Invitrogen) or the polyclonal rabbit antibodies anti-FBN1, anti-FBN2, or anti-fibronectin (pAB 2033, MilliporeSigma, Burlington, MA) (36, 54–56).

## Author contributions

DH designed and performed the experiments, analyzed the data and wrote the manuscript. SSA designed experiments, analyzed the data and wrote the manuscript. ST analyzed the *Adamtsl2*-Prx Achilles tendon dimensions. SMA and DEB performed transmission electron microscopy. RS provided the *Scx*-Cre deleter strain. All authors edited and approved the manuscript.

## Acknowledgements

This work was supported by the National Institutes of Health (awards AR070748 to D.H., AR53890 to S. A, and AR44745 to D. B.). We thank Dr. Dieter Reinhardt (McGill University, Montreal, Canada) for providing the fibrillin-1 antibody and Dr. Robert Mecham (Washington University, St. Louis, USA) for providing the fibrillin-2 antibody. We thank Dr. Judith Drazba from the Digital Imaging Core (Cleveland Clinic Lerner Research Institute) for help with two photon imaging. This work utilized a Leica SP5 confocal/multi-photon microscope that was purchased with partial funding from National Institutes of Health SIG grant 1S10RR026820-01.

## Non-standard Abbreviations

ADAMTSL: a disintegrin-like and metalloproteinase domain with thrombospondin-type 1 motif-like
DMEM: Dulbecco’s Modified Eagle Medium
ECM: extracellular matrix
FBN1: fibrillin-1
GD: geleophysic dysplasia
KOMP: Knockout Mouse Project
MFS: Marfan syndrome
PBS: phosphate-buffered saline
TGFβ: transforming growth factor-β
WMS: Weill Marchesani syndrome
RT-qPCR: real-time quantitative PCR

---

**Supplemental Figure 1.**
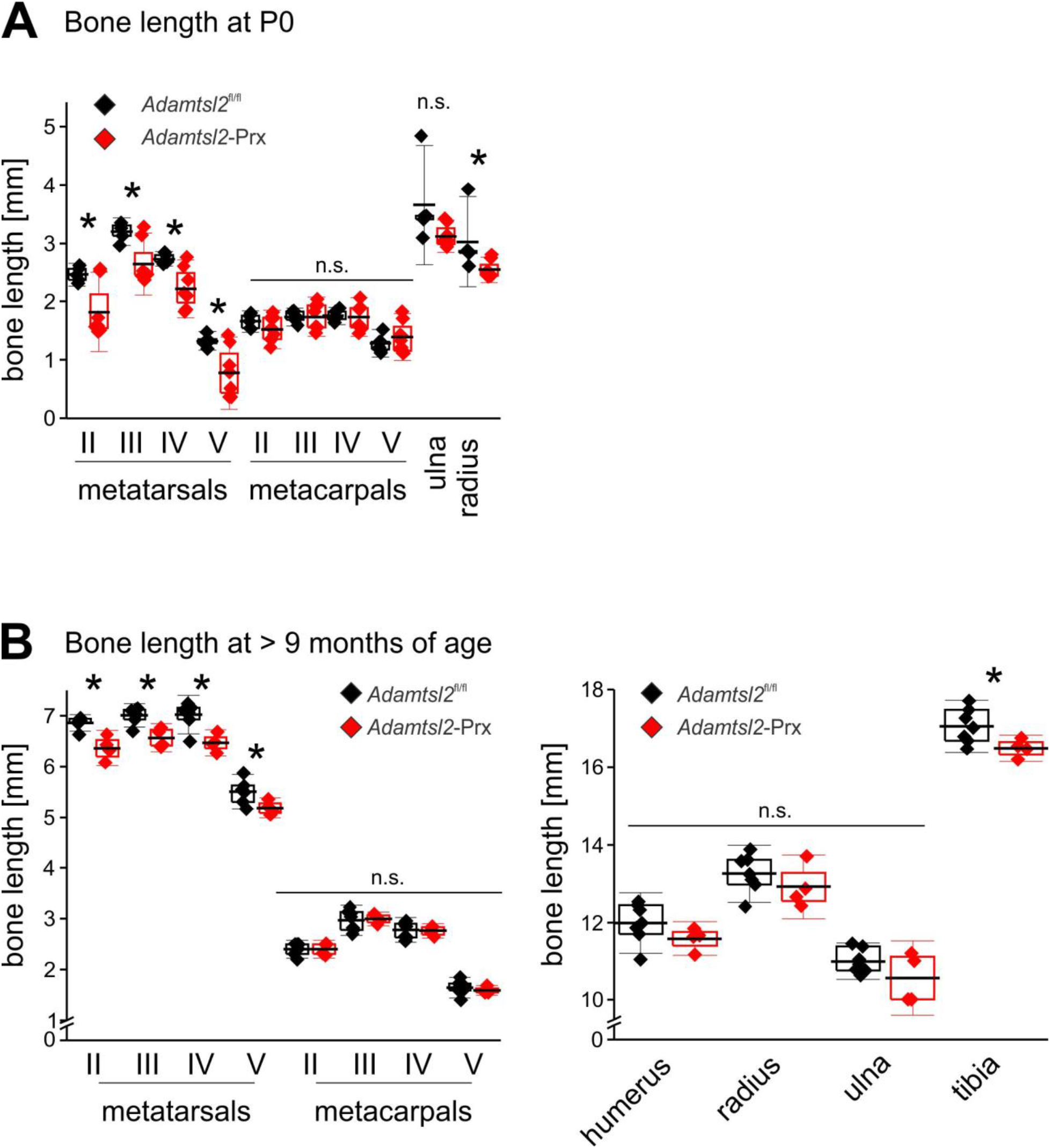
Bone length alterations in newborn and skeletally mature (>9 month old) mice. **(A)** Measured lengths of newborn metatarsals, metacarpals, ulna and radius demonstrate relative shortening of metatarsals and radius in *Adamtsl2*-Prx limbs (n=5-8). **(B)** Measured lengths of skeletally mature metatarsals, metacarpals, ulna and radius demonstrates persistence of bone shortening in metatarsals and the tibia in *Adamtsl2*-Prx limbs (n=4-7). P-values were calculated with a 2-sided Student t-test. * p<0.05; n.s., not significant. The box indicates the 25^th^-75^th^ percentile, the line indicates the mean and whiskers indicate standard deviation.

**Supplemental Figure 2.**
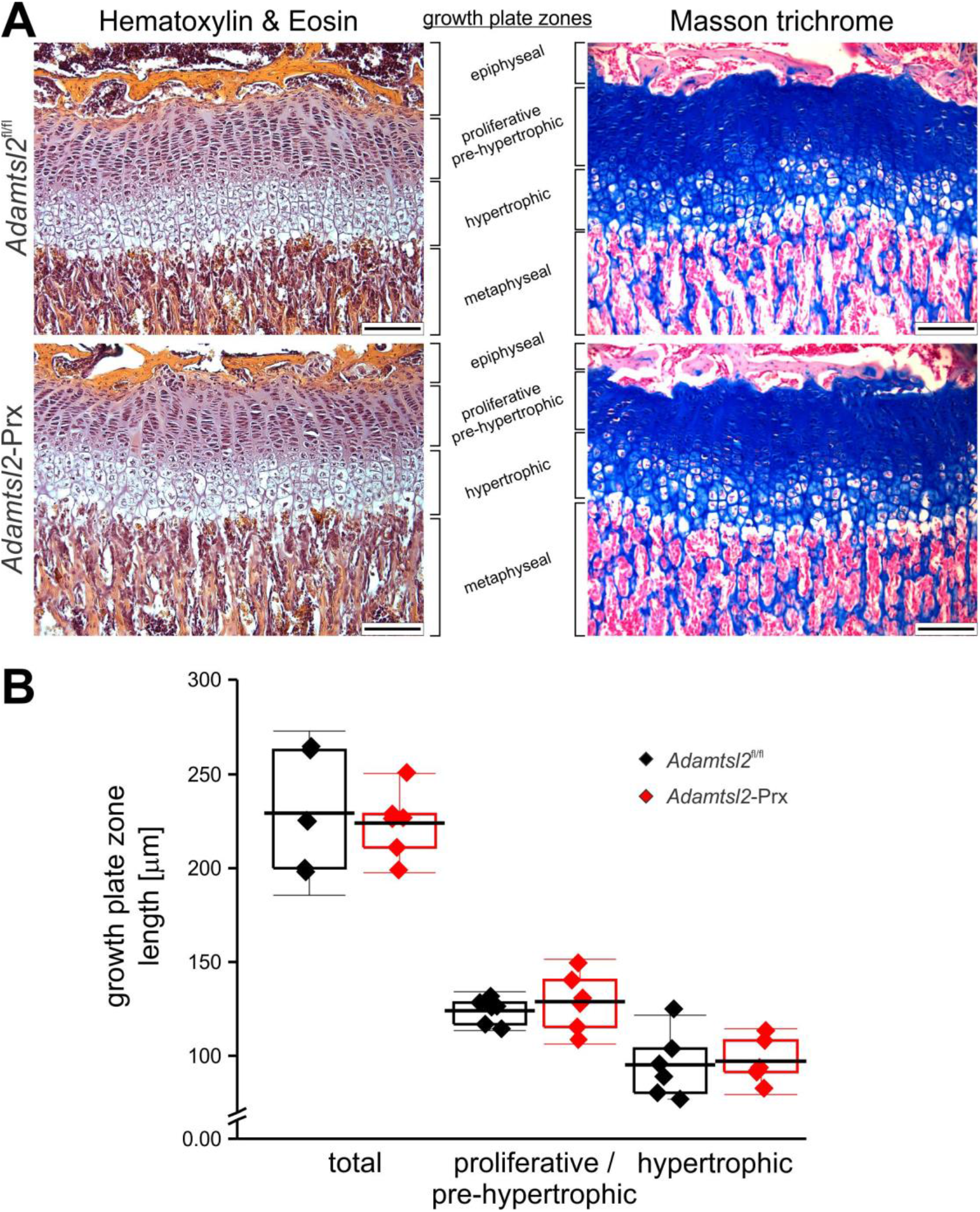
Normal growth plate architecture and morphometry in *Adamtsl2*-Prx limbs. **(A)** Hematoxylin and eosin (left panels) and Masson Trichrome (collagen, blue, right panels) stained growth plates of proximal tibiae indicate comparable histological organization of the growth plates. Images shown are representative of n=6 knees from n=3 littermates per genotype. **(B)** Quantification of overall growth plate thickness, proliferative / pre-hypertrophic, and hypertrophic zones identified no differences in growth plate metrics in *Adamtsl2*-Prx tibia compared to control (n=3). P-values were calculated with a 2-sided Student t-test. The box indicates the 25^th^-75^th^ percentile, the line indicates the mean and whiskers indicate standard deviation.

**Supplemental Figure 3.**
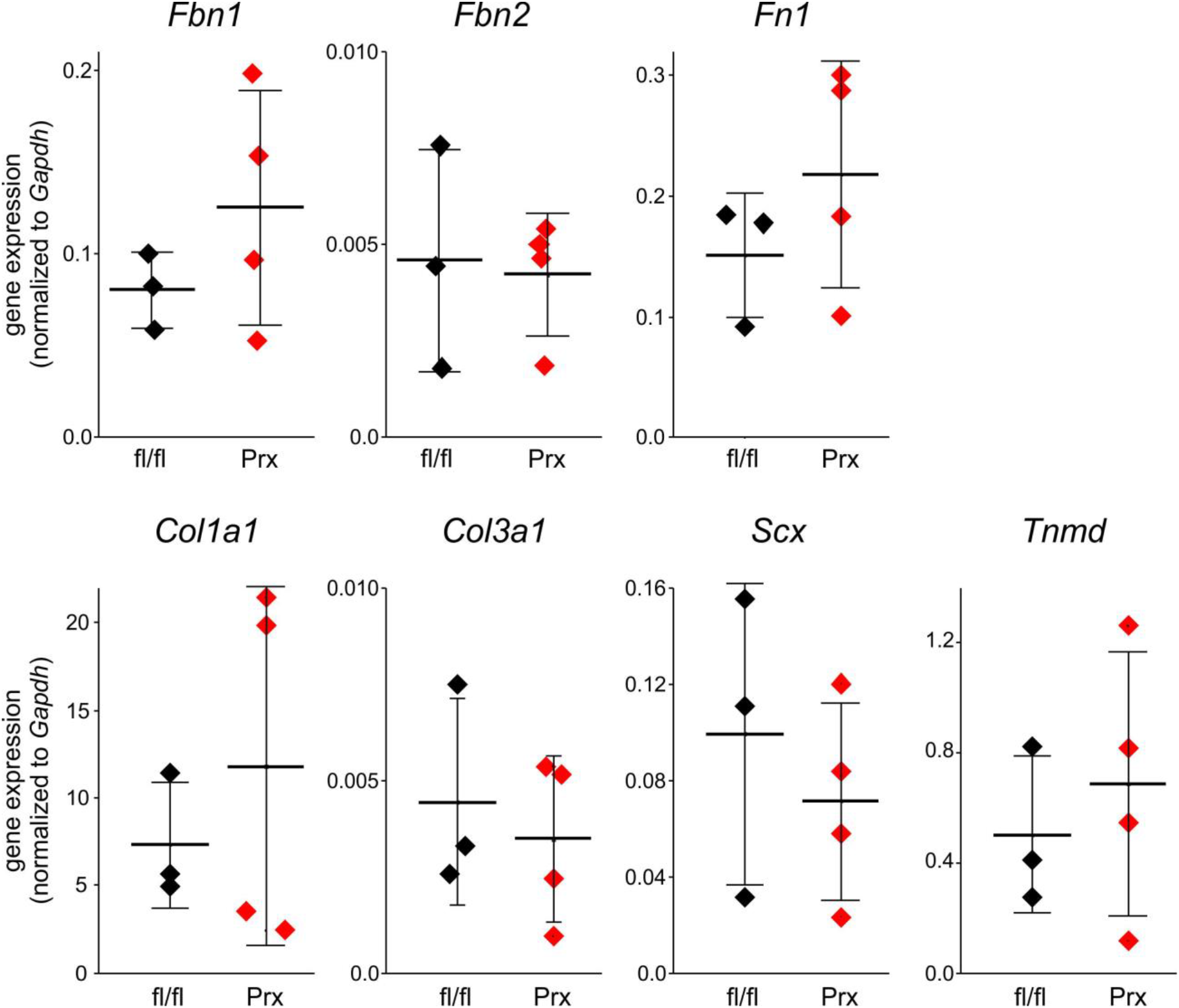
Quantitative real-time RT-PCR analysis of relevant ECM and tendon transcripts showing comparable levels to control (fl/fl) in *Adamtsl2*-Prx (Prx) Achilles tendon. Genes in the following categories were analyzed: <u>ECM:</u> Fibrillin-1 (*Fbn1*), −2 (*Fbn2*), fibronectin (Fn1); collagen I (*Col1a1*), collagen III (*Col3a1*); <u>Tenocyte markers:</u> Scleraxis (Scx), tenomodulin (*Tnmd*) (n=3, 4). P-values were calculated with a 2-sided Student t-test. The line indicates the mean, and whiskers indicate standard deviation.

